# Self-organized Regulation of Group Size and Number in Natural and Artificial Collectives

**DOI:** 10.64898/2026.08.25.746978

**Authors:** Tianfu Zhang, Suet Lee, Heiko Hamann

## Abstract

From animal societies to self-organizing multi-agent systems, collectives adapt their group structure to tasks and environments. However, how they determine appropriate group sizes and the number of subgroups to form remains unclear. We formulate the Group Size and Number Regulation Problem (GSNRP), which asks how individuals regulate group sizes and numbers using only local information. In a first step, we establish a graph-theoretic model demonstrating that simple following behavior suffices to form group structures that match theoretical expectations, but is insufficient for active regulation of group size and number. In a second step, we operationalize individual group-size preferences in a decentralized fission–fusion mechanism based on perceived group size and an internal size preference, with no runtime access to population size or total group number. Through multi-agent simulations, we validate that this mechanism achieves stable convergence across three signaling regimes, from position-only sensing to continuous group-size communication. Using tracking data from wild white-nosed coatis (mammals in the raccoon family), we calibrate individual group-size preferences and show that the controller recovers selected group-size, subgroup-count, and transition statistics. This in-sample case study demonstrates descriptive consistency with natural fission–fusion dynamics, but does not establish the underlying behavioral mechanism. These results suggest that natural and engineered collectives may share local principles of perception, preference, and response for regulating group structure.

## 1 Introduction

In natural systems, the ability of individuals to organize into (dynamic) groups is essential to coordinate, adapt, and scale. Such grouping allows individuals to reduce risk, share information, and exploit resources more effectively, while flexibly balancing the costs and benefits of collective living. When viewed through an engineering lens, these biological grouping mechanisms can be interpreted as localized interaction rules, and understanding how such rules give rise to dynamic collectives in nature can provide important insights into the design of robust, adaptive multi-agent systems.

The dynamic organization of groups is fundamentally influenced by effects of group size on the individual animal on many timescales: evolutionary, ecological, and behavioral. Changes in group size are commonly accompanied by density-dependent effects on survival. As early as Allee’s work on animal aggregations, it was observed that survival or reproductive success can depend on local population density or group size, particularly at low densities Allee (1927); Pride (2005); Courchamp et al. (2008). For example, through intraspecific cooperation, individuals can reduce the probability of predation by aggregating. According to the selfish herd hypothesis, individuals at the center of the group have a survival advantage Hamilton (1971). When group sizes exceed a certain threshold, negative effects, such as competition for resources, social interference, and disease transmission, come to outweigh the benefits of group living and lead to a decrease in individual fitness (e.g., observed as a negative per capita population growth rate). Therefore, the relationship between density and per-capita growth rate has been described as an inverted

U-shaped curve Allee (1927) and similarly individual fitness over group size Sibly (1983). The Allee effect describes inverse density dependence at low density, assuming a critical density (or critical group size) below which reduced aggregation impairs survival or reproduction Courchamp et al. (1999). Hence, there is an intermediate optimal population density, which we understand as an optimal group size Brown (1982); Markham et al. (2015). At this size, the group can collaborate effectively to realize synergistic benefits while avoiding excessive competition for shared resources.

If individuals are free to join or leave groups (e.g., not forced to leave), fission–fusion dynamics may allow them to track group sizes that maximize their expected individual fitness. This strategic behavior, in turn, would ignore optimal group size, because a solitary individual can still gain by joining a group already at its optimum. The optimum is therefore unstable and difficult to maintain collectively in the resulting group dynamics Sibly (1983). Environmental conditions, resource availability, and social relationships typically influence the behavior of social animals.

Individuals may choose to leave or join a group in the short-term. The accumulation of such behavioral decisions based on local perceptions results in the macro-level fission-fusion group structure Aureli et al. (2008). Fission–fusion dynamics have accordingly been framed as collective decision processes in which individual needs, social relationships, and the scale of information flow jointly influence whether groups remain cohesive, split, or merge Sueur et al. (2011). This dynamic organization allows animals to respond flexibly to environmental uncertainty, temporarily reduce competition within the group, and improve individual survival by balancing the benefits of sociality with the costs of group living. The evolutionary adaptability of this dynamic organization Sumpter (2006) suggests that local, perception-based decisions, such as joining or leaving groups, constitute a behavioral mechanism with significant advantages.

In engineering systems, similar problems of group formation and dynamics arise in the context of task allocation Brutschy et al. (2014); Debie et al. (2023) and congestion mitigation Kuckling et al. (2024). For example, in multi-robot task execution, individual agents form groups of different numbers and sizes according to task requirements and environmental conditions. The size and number of groups directly affect the system’s operational efficiency. Groups may be too small to complete a collaborative task or too large, leading to conflicts over access to shared resources, such as communication channels or physical spaces Hamann and Reina (2021). For example, in large-scale multi-robot systems, system performance exhibits a nonlinear response to increasing robot density Østergaard et al. (2001), and sometimes even a sudden transition from efficient operation to a fully congested state, resulting in two phases of system behavior Kuckling et al. (2024).

Many models of group dynamics have adopted a macroscopic, phenomenological perspective, focusing on population-level quantities such as group size, density, or per-capita growth rates.

These approaches have been successful in characterizing general patterns and identifying conditions under which positive or negative density-dependent effects arise, often without requiring the specification of the detailed mechanisms at the individual level. At the same time, such descriptions leave open the question of how local interactions and general behavioral constraints among individuals generate the observed collective dynamics. Investigating microscopic interaction mechanisms can thus complement macroscopic models by probing their mechanistic plausibility.

This perspective is also relevant to engineered multi-agent systems, where collective behavior must emerge from explicitly defined local rules and comparable tradeoffs between cooperation and interference. In this paper, we study a key question (illustrated conceptually in Fig. 1): Can individuals in either animal or engineered collectives self-organize to regulate both the size and number of groups in response to changing environments or task demands using only local behavioral mechanisms?

**Figure 1:**
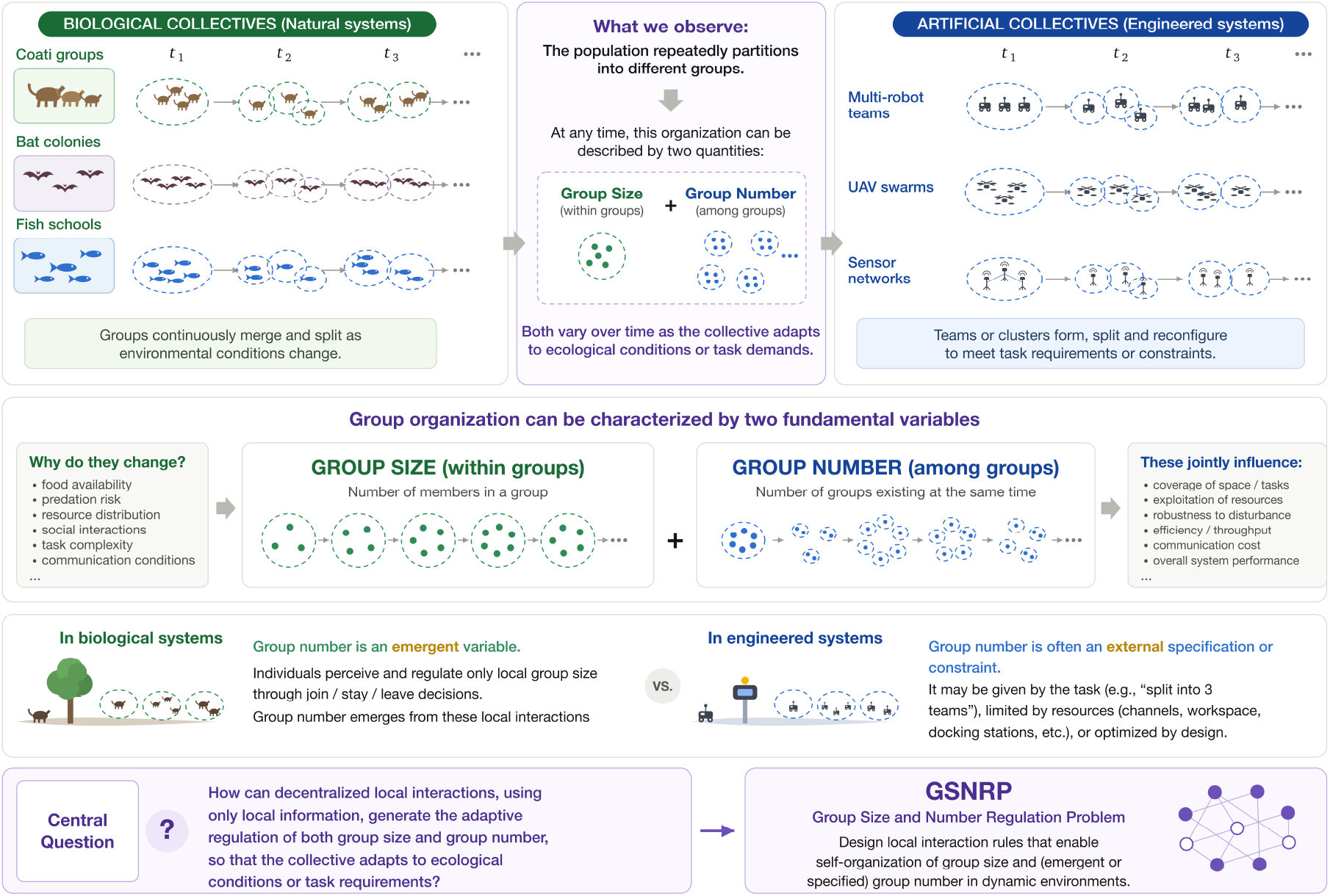
Group size and group number as two complementary descriptors of collective organization in biological and engineered systems, motivating the Group Size and Number Regulation Problem.

Complementary individual- and agent-based models have shown that fission–fusion organization can emerge from simple foraging in heterogeneous environments Ramos-Fernández et al. (2006), the dynamics of grooming and social networks Sueur et al. (2011); Sueur and Maire (2014), individual roost-selection decisions Perony et al. (2022), and heterogeneous group-size preferences Guerra et al. (2020). We study whether individual group-size preferences can be operationalized as a decentralized feedback mechanism that jointly regulates group size and group number under restricted information.

The combined and simultaneous regulation of both group size and the number of groups is key in natural and engineered systems. In this paper, we pose the problem we call the *Group Size and Number Regulation Problem (GSNRP)* and construct a decentralized behavioral model inspired by biological systems (see Fig. 2 for an overview). We propose a general framework and provide a formal modeling approach that help to understand the mechanisms shaping fission-fusion behavior in animal populations. We validate the feasibility of our model in engineered systems through theoretical analysis and multi-agent simulations. We also compare the group structures generated by the model with tracking data from wild animals in a calibrated empirical case study. This comparison assesses descriptive consistency across selected group-size, group-number, and transition statistics, without aiming to identify the behavioral mechanism operating in the animals.

**Figure 2:**
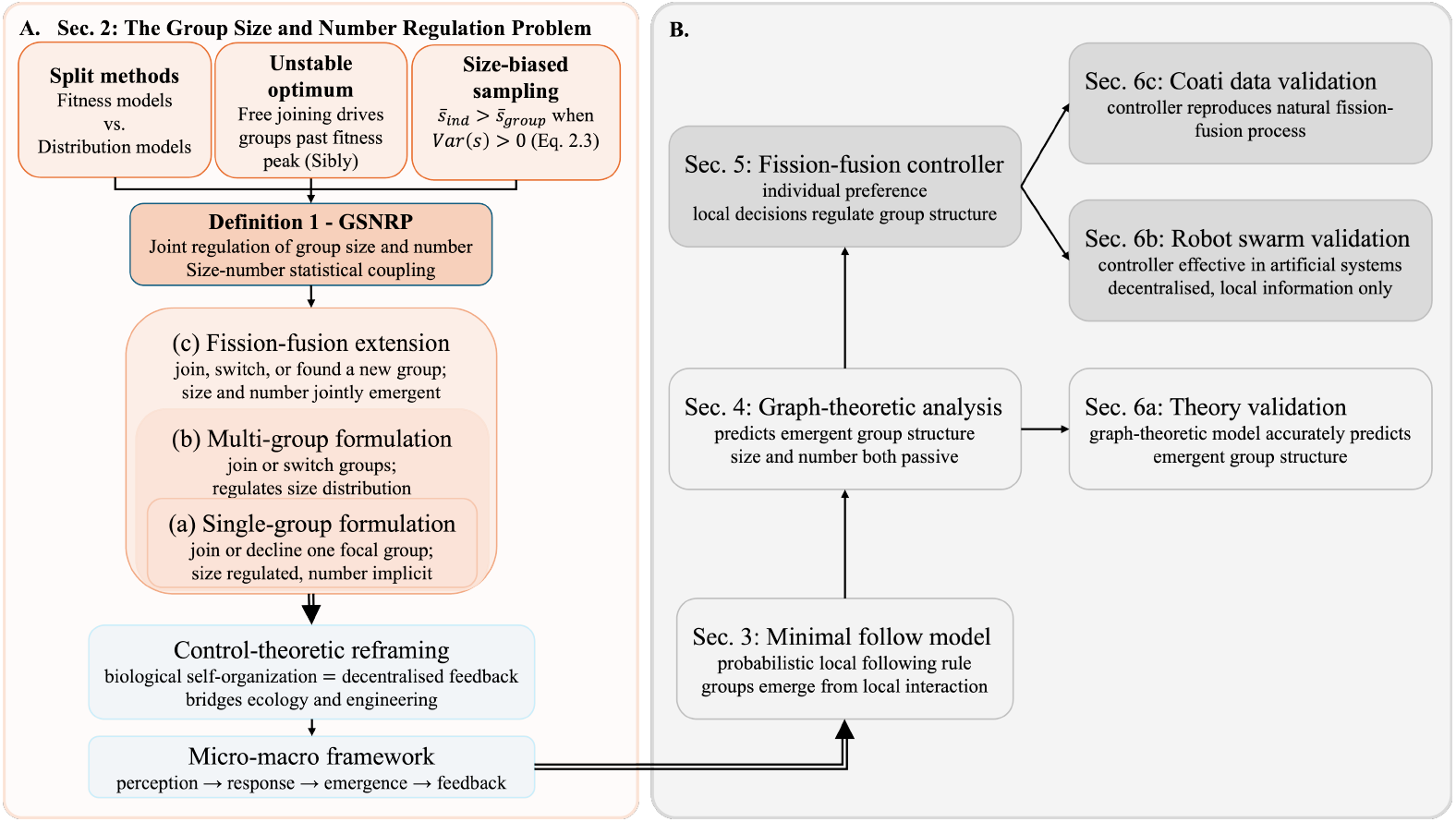
Structural overview of the paper. (A) Three motivating observations converge on Definition 1 of the Group Size and Number Regulation Problem (GSNRP). Three nested formulations of increasing generality are introduced: single-group sequential joining 2.1, multi-group choice 2.2, and fully endogenous fission–fusion dynamics 2.3. Together, these formulations provide the basis of a micro-macro framework for biological self-organization and decentralized control and connect to the modeling pipeline. (B) Starting from the micro–macro framework, the pipeline proceeds from a minimal local following model 3 through graph-theoretic analysis 7.1 and a decentralized fission–fusion controller 4, validated in theoretical 5.1, engineering 5.2, and biological 5.3 settings.

With this work, we seek a common conceptual framework for describing how natural and artificial systems regulate their group structure through similar local mechanisms of perception, preference, and response.

## 2 The Group Size and Number Regulation Problem

Existing work has examined optimal group size, individual joining and leaving decisions, fission–fusion dynamics, and stationary group-size distributions from complementary perspectives Sibly (1983); Allee (1927); Krause and Ruxton (2002). These approaches typically emphasize either the micro level, by analyzing how individual decision-making and fitness shape group membership, or the macro level, by characterizing the resulting population-wide distribution of group sizes. In a population of fixed size, however, these two perspectives are intrinsically coupled: local decisions determine the evolving partition of individuals into groups, while that partition simultaneously determines both the distribution of group sizes and the total number of groups. Here, we focus on the question whether group size and group number can be jointly regulated in a self-organizing way? And we ask the more specific question: how can locally interacting individuals use perceived group size and internal size preferences to regulate an evolving group partition through decentralized feedback, without global information? We argue that group size and group number must be studied jointly as a micro–macro consistency problem, linking individual-level decision mechanisms to emergent population-level group-size distributions.

From a theoretical perspective, the assumption of an optimal group size implies that individual fitness is an inverted-U-shaped function (concave, unimodal) over group size, with an optimum that balances cooperation benefits and competition costs. If individuals are free to join larger groups due to their individual benefits, this will drive groups beyond the fitness optimum Sibly (1983).

Even when an optimal group size exists with respect to individual fitness, it is generally not a stable equilibrium under free-joining dynamics.

Also, the animal group-size distribution is often heavy-tailed or power-law distributed Bonabeau et al. (1999); Niwa (2003), meaning that small groups are more frequent. This is known as the size-biased sampling effect (typical group size effect). Heavy-tailed group-size distributions cause individuals to experience groups that are systematically larger than suggested by the group-level mean. Let *P*_grp_(*s*) denote the group-centered distribution, that is, the probability that a uniformly randomly sampled group has size *s*, and let *P*_ind_(*s*) denote the individual-centered (experienced) distribution, that is, the probability that a uniformly randomly sampled individual belongs to a group of size *s*. Then we get

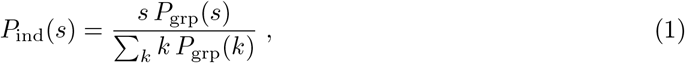

that is, large groups are disproportionately more likely from the individual perspective and are generally larger than the population-level mean (Σ_*s*_ *s P*_ind_(*s*) *>* Σ_*s*_ *s P*_grp_(*s*)) Jovani and Mavor (2011); Reiczigel et al. (2008). The representative group size one reports depends on whether one adopts a group-centered or an individual-centered perspective.

In summary, and taking a modeling perspective, group size and group number should not be treated as independent outcomes. Instead, they should be understood as jointly regulated variables in a dynamically coupled system. The problem is therefore twofold: first, whether a decentralized mechanism for jointly regulating group size and number can exist at all under local information only; and second, how such a mechanism operates so that individual-level incentives and population-level statistical structure remain compatible.

We formalize this challenge as the Group Size and Number Regulation Problem (GSNRP).

More specifically, we ask how a group of locally interacting individuals, without global information, self-organizes into a dynamic group configuration in which both the distribution of group sizes and the number of groups are adaptively regulated. By coupling size and number regulation, GSNRP bridges the gap between fitness-based optimality arguments and empirically observed heavy-tailed group structures, providing a more complete perspective for biological fission-fusion societies and artificial multi-agent systems.

### Definition 1

(Size–Number Statistical Coupling and GSNRP). Consider a system with *N* individuals partitioned into *G* groups with sizes given as set 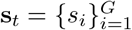, where *s*_*i*_ ∈ ℕ and 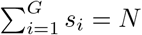. Define the *group-level mean size* and the *individual-level experienced size* as

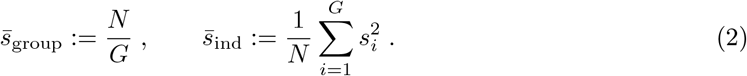

Equivalently, 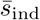 is the expectation of group size under the size-biased distribution experienced by a uniformly sampled individual. Using the sample-based variance 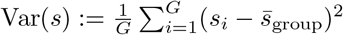, we get

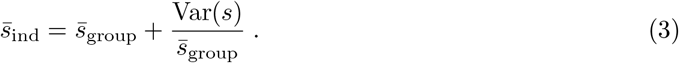

This makes the earlier inequality Σ_*s*_ *s P*_ind_(*s*) *>* Σ_*s*_ *s P*_grp_(*s*) explicit: the individual-level mean exceeds the group-level mean by exactly 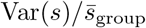, so the gap is governed entirely by the variance of the size distribution. Hence, whenever the size distribution has a nonzero variance (Var(*s*) *>* 0), it holds that 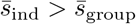, implying that the group count *G* or the mean group size 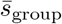 alone cannot characterize the size actually experienced by individuals. For example, both {99, 1} and {55, 45} give 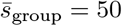, yet 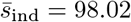 and 50.5 respectively, so they are identical at the group level but nearly a factor of two apart at the individual level.

This mathematical observation highlights the need for joint regulation of group size and group number, motivating our central question. We formalize this central question as the *GSNRP* : under the conservation constraint 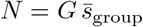, does there exist a decentralized mechanism that jointly adjusts the group-size distribution {*s*_*i*_} and the group number *G* so as to form a stable group structure? Here, stability is statistically significant: despite ongoing fission-fusion dynamics, the distributions of group numbers and group sizes remain stable, while the distribution of group sizes experienced by individuals, resulting from size-biased sampling, also remains stable over time.

Next, we introduce three increasingly rich formulations of group formation, ranging from binary sequential acceptance within a focal group to choice among multiple coexisting groups to fully endogenous fusion–fission dynamics with both arrivals and departures. These three formulations are nested: Sec. 2.1 and Sec. 2.2 characterize the restricted settings addressed by existing work, and only the fission–fusion formulation 2.3 is instantiated in the controller of Sec. 4 and evaluated in Sec. 5.2 and 5.3.

### 2.1 Sequential Single-Group Formulation

An important aspect of group-size choice models is that they implicitly assume an *iterated sequential decision-making* process, as already suggested by Sibly Sibly (1983) and related approaches. Consider a population of *N* individuals arriving sequentially at a focal group. Let *s*_*t*_ ∈ ℕ denote the group size observed by the *t*-th individual at decision time *t*. Each individual makes a one-shot binary decision

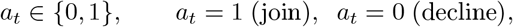

based on a utility function *U* (*s*_*t*_). Decisions are irreversible, and group size evolves as

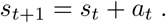

This process repeats across many individuals until convergence to a fixed point *s*^*^ := lim_*t*→∞_ *s*_*t*_ satisfying

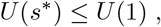

so that no further individual prefers to join (Fig. 3E). Equilibrium group size thus emerges as the fixed point of sequential acceptance decisions. Notably, Sibly-type models regulate size but leave group number implicit, thereby addressing only a restricted version of GSNRP.

**Figure 3:**
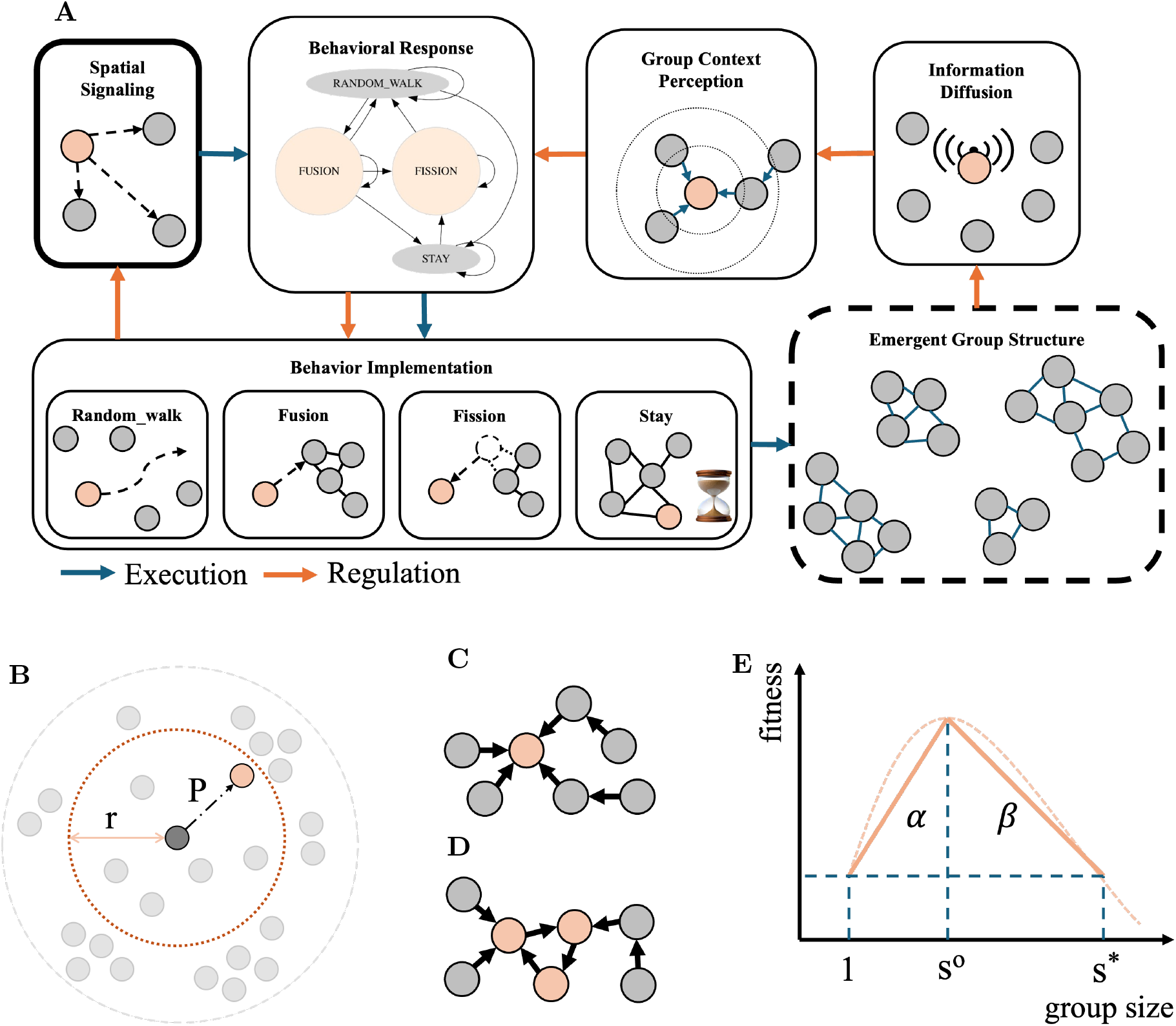
(A) Overview of the decentralized group regulation framework. The framework addresses the problem of group structure regulation, encompassing the control of group size, number, composition, and spatial/relational organization. Blue arrows indicate the execution flow, where selected behaviors are implemented, leading to the structure of the overall group. Orange arrows represent the regulation flow, in which agents evaluate their local group context to decide on behavioral responses. (B) Minimal behavioral model. Each agent (gray node) perceives neighbors within its perception radius (gray circle) and decides with probability P whether to follow a target within the follow range *r* (orange dashed circle). The focal agent (central gray) thus probabilistically connects to a neighbor (orange) within the local field, forming a directed connection. Two types of agent group structures: (C) Directed tree (D) Directed pseudotree with cycle. (E) Fitness curve simplified using slope parameters *α* and *β*.

### 2.2 Sequential Multi-Group Formulation

To move beyond the focal-group setting of Sibly-type models, we next consider a sequential multi-group formulation Gueron and Levin (1995) in which individuals do not merely decide whether to join a single encountered group, but instead choose among several coexisting groups. We now consider *G*_*t*_ coexisting groups at time *t* with sizes

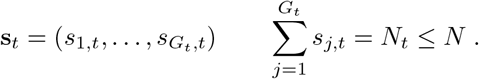

Each arriving individual chooses among groups according to

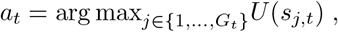

so individuals effectively select from a set of group sizes rather than responding to a single focal group. The chosen group evolves as

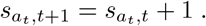

The system converges when no arriving individual prefers any alternative group under *U* (·), yielding a distributional equilibrium across groups.

### 2.3 Fission–Fusion Extension

Finally, we allow endogenous departures (individual-initiated exits) in addition to arrivals. At time

*t*, an individual currently in group *i* of size *s*_*i,t*_ chooses among

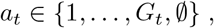

where *a*_*t*_ = *j* denotes moving to group *j* (including *j* = *i*, i.e., staying) and *a*_*t*_ = ∅ denotes leaving to found a singleton group. Decisions follow

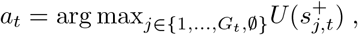

with

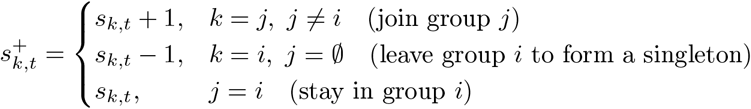

This formulation makes explicit that leaving can be interpreted either as selecting again from the available group sizes or as forming a singleton group. Does there exist a decentralized rule under which both *G*_*t*_ and the group-size frequency distribution (GSFD) converge to a stable joint distribution? This extension reveals GSNRP as a fully endogenous partitioning problem, in which both group size and the number of groups arise as emergent outcomes of decentralized mobility decisions.

#### Control-theoretic perspective

The GSNRP can be interpreted as a decentralized regulation problem with respect to emergent macroscopic variables. Let the system state at time *t* be characterized by the group configuration (*G*_*t*_, **s**_*t*_), where *G*_*t*_ denotes the number of groups and 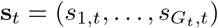 their sizes. Individual decisions *a*_*t*_, taken using only local information, act as microscopic control inputs that indirectly shape these macroscopic variables through their aggregate effect. Thus, (*G*_*t*_, **s**_*t*_) are not directly controlled but emerge from the closed-loop interaction between decentralized decision rules and the conservation constraint 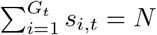.

From this perspective, GSNRP asks whether there exist decentralized policies such that the induced stochastic process over group configurations stabilizes (in an appropriate sense) around a desired regime for the number of groups and the size distribution.

### 2.4 Micro-macro regulation framework

The control-theoretic interpretation above provides a unifying perspective that links biological grouping behavior to decentralized regulation principles. As summarized in Fig. 3A, local perception and behavioral responses observed in biological systems can be interpreted as microscopic control mechanisms acting on emergent collective variables. Individuals perceive local group conditions *s*_*j,t*_ and respond through elementary fission–fusion decisions *a*_*t*_, such as joining, staying, or separating. These microscopic decisions accumulate into macroscopic collective variables **s**, including group size, group number, and spatial organization. The resulting group structure then feeds back onto subsequent individual decisions, forming a closed-loop regulation process between microscopic behavior and macroscopic organization. From this perspective, GSNRP is both a biological self-organization problem and a decentralized control problem.

## 3 Minimal interaction model for emergent group formation

To address the GSNRP, we begin by examining how group structures can emerge from simple local interactions among individuals. In many biological groups, individuals often exhibit behaviors of following and/or aligning with neighboring members Reynolds (1987); Vicsek et al. (1995). For example, in schools of fish Couzin et al. (2002), flocks of birds Cavagna et al. (2010), or groups of mammals Ginelli et al. (2015), individuals typically adjust their behavior based on the positions or movement directions of their neighbors. Such local interactions based on partial information only can still result in stable group structures.

In this section, we abstract the widely observed following behavior into a minimal interaction model. This model captures the simplest local rule that allows groups to emerge and serves as a baseline for understanding the relationship between individual behavior and collective structure.

### 3.1 Local sensing and following

We consider a system of *N* homogeneous individuals distributed in a two-dimensional space. Let *V* denote the set of all individuals. Each individual can sense only neighboring individuals within a range determined by the sensing radius *r*, which may correspond biologically to perceptual constraints such as visual, acoustic, or social sensing limits Sueur et al. (2011). We define the neighborhood of an individual *i* ∈ *V* at time *t* as

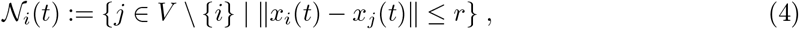

where *x*_*i*_(*t*) ∈ ℝ^2^ denotes the position of individual *i* at time *t*, and ∥·∥ is the Euclidean norm.

Each individual *i* adopts a simple probabilistic rule: with probability *P*, it follows a uniformly randomly chosen neighbor from *N*_*i*_, and with probability 1 − *P*, it acts independently. Following time *t*, individual behavior is influenced by the information of neighboring individuals and the individual they follow, if any. The individual they follow is called the target. We represent the following as a map

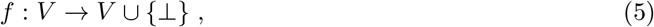

where *f* (*i*) = *j* means individual *i* follows individual *j*, while *f* (*i*) = ⊥ means that individual *i* does not follow any individual.

We analyze this model from a graph-theoretic perspective (see Appendix 7.1). Since each individual follows at most one target, the resulting follow graph is a directed pseudoforest: every connected component is a group that either has a root following nobody (a directed tree, Fig. 3C) or contains a single directed cycle that all remaining members eventually reach (Fig. 3D). Two dimensionless quantities govern this structure. The geometric density *ϕ* := *λπr*^2^, with population density *λ* = *N/A*, is the expected number of neighbors within interaction range and sets the probability *ρ* ≈ 1 − *e*^−*ϕ*^ that an individual has any neighbor at all. Combined with the follow probability *P*, this gives the effective following probability *η* := *ρP*, that is, the probability that an individual actually establishes a following edge. The expected number of groups is then

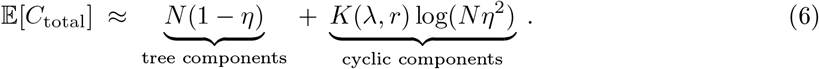

The factor *K*(*λ, r*) is calibrated on a small subset of conditions. We test eq. (6) against simulations in Sec. 5.1.

### 3.2 Limitations of pure following behavior

This minimal model based on local following behavior has the potential to predict the group structure. The predicted group structure is a deterministic function of *λ, r*, and *P*. The model answers a descriptive question: which patterns arise under given parameters. It does not answer the regulatory question posed by GSNRP: how individuals maintain group size near a desired target. Individuals respond only to local neighbor relations. They receive no signal about whether their group is too small, too large, or about right. Group size is then a passive consequence of environmental parameters, not an actively stabilized quantity. Closing this gap requires individuals to perceive the group state and respond to it, which is the step we take next.

## 4 Subjective group size preferences: a behavioral strategy for solving GSNRP

In social animals, group size plays a critical role in shaping individual fitness. Numerous studies Markham et al. (2015); Brown (1982) in behavioral ecology have suggested a nonlinear, inverted-U-shaped relationship between fitness and group size: small groups may suffer from insufficient collaboration and higher predation risk, and potentially oversized groups may experience increased competition. However, achieving such adaptive group sizes is often difficult to maintain solely through random processes. This suggests that individuals may be required to perceive and respond to group size Agrillo and Bisazza (2018); Mukherjee and Bassler (2019).

Whether in biology, where individuals perceive and respond to the current state of their group, or in engineering, where feedback signals drive a system toward a desired setpoint Åström and Murray (2021), a common principle emerges: individual behavior is governed by the discrepancy between the actual group state and an internal reference. Motivated by empirical evidence for group-size choice Pritchard et al. (2001) and by previous preference-driven fission–fusion

models Guerra et al. (2020), we operationalize subjective group-size preference as a decentralized feedback variable: each individual *i* possesses a fixed internal desired group size 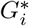 and adjusts its behavior accordingly Pritchard et al. (2001). We emphasize that 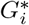 is an individual trait rather than a description of the collective. Each agent holds a fixed internal scalar estimated from population-level observations during calibration, as is standard practice for any agent-based model parameter, but carries no runtime information about the current population size *N* or group count *G*. If *N* were changed at runtime, any agent’s behavior would remain unchanged.

To test this hypothesis, we implement a multi-agent conceptual framework (Fig. 3A). We propose a decentralized control strategy in which each individual only has information about the current group size. Each individual compares its current group size with an internal desired size and accordingly initiates an individual fission or fusion behavior. This individual-level regulation mechanism is integrated into the coordination framework introduced in Sec. 2.4. We aim to examine whether such local behavioral rules are sufficient to produce stable and self-regulating group structures at the collective level, thereby addressing the GSNRP.

### 4.1 Behavioral state-machine formulation

To operationalize subjective group size preference, we translate it into a simple behavioral state machine. The key idea is that individuals do not continuously optimize an explicit objective function. Instead, we switch to an agent-based model, in which each individual switches among a small number of discrete behaviors depending on how its current group size compares with its internal target. This yields a decentralized control mechanism that is simple enough to analyze and implement, yet rich enough to generate adaptive group-level dynamics.

The core of the proposed strategy consists of four discrete behavioral states: `Random_Walk`, `Fusion`, `Fission`, and `Stay` (Fig. 4). Together, these states define a minimal repertoire of actions through which an individual can contribute to the formation, maintenance, and restructuring of groups.

**Figure 4:**
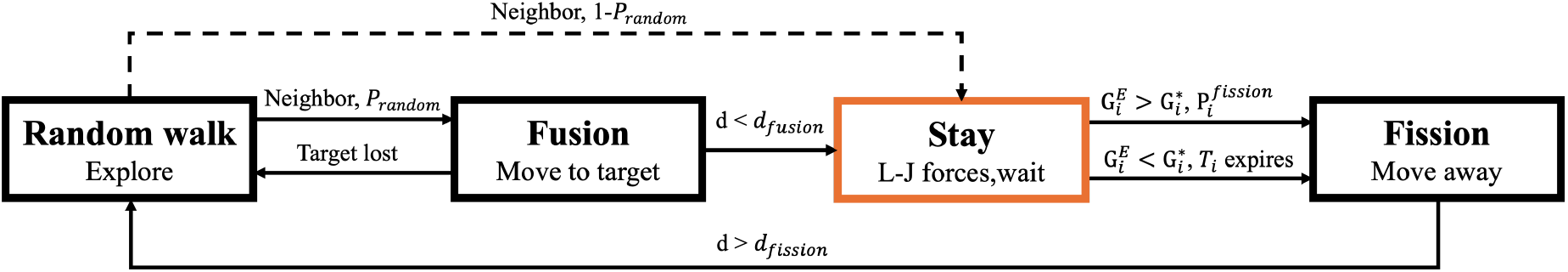
Behavioral state machine of the decentralized fission-fusion controller

The state `Random_Walk` represents exploratory behavior in the absence of a satisfactory local grouping condition. `Random_Walk` provides the mechanism by which isolated individuals search for aggregation opportunities. The state `Fusion` is activated when an individual in `Random_Walk` detects a target within its follow range. The individual then abandons exploration and moves toward the detected target in order to join or form a group. The state `Fission` is activated when the current group is perceived as too large. The individual then initiates behavior that increases separation. The state `Stay` represents a locally satisfactory condition. When the current group size is sufficiently close to the individual’s internal preference, no corrective action is taken.

Behavioral regulation emerges from the transition logic among these four states, the definitions of the states, and the transition conditions (see Appendix 7.2). The behavioral state determines the motion policy that governs the agent’s trajectory. At each time step, agents update their position according to state-dependent movement rules, including stochastic exploration, target-directed motion, separation behavior, or local attraction–repulsion interactions.

## 5 Experiments

In the following, we progressively demonstrate the efficacy of the proposed framework for regulating group size and number, proceeding sequentially from theoretical validation to algorithmic evaluation, and then to simulations and real-animal data.

### 5.1 Predictability of group number

To evaluate the predictive performance of the proposed theoretical model (see Sec. 3 and Appendix 7.1), we compare its predictions with simulation averages over a large parameter sweep. Figure 5A presents the comparison of the number of components as a function of the composite factor *η* = *ρP*, which represents the effective probability that an individual forms a following connection. A following edge can only emerge when an individual both encounters at least one neighbor within interaction range (captured by *ρ*) and decides to follow (captured by *P*).

**Figure 5:**
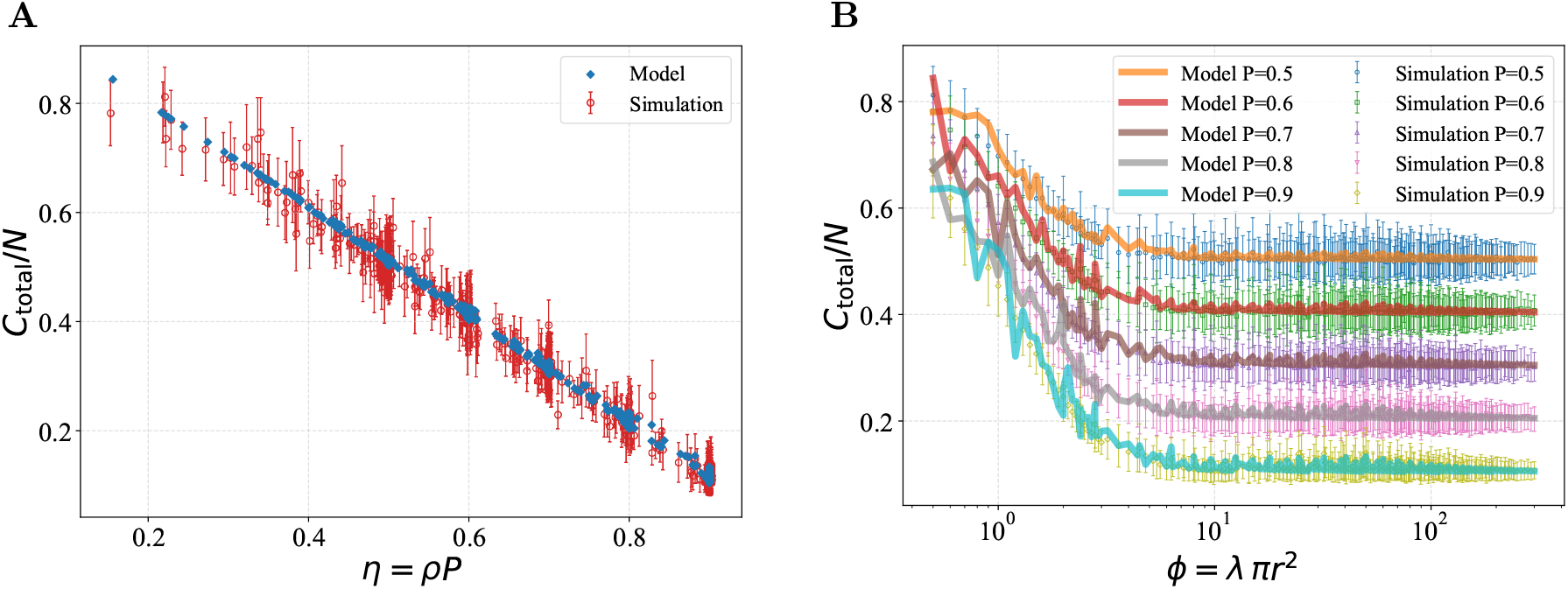
Comparison of the number of components under different parameters: (A) Comparison of the number of components as a function of the composite factor *η* = *ρP*, which captures the coupled effect of the follow probability and spatial distribution. Both theoretical predictions and simulation results show a decreasing trend as *η* increases, and the two remain highly consistent, validating the proposed model’s predictive capability at the average level. (B) Variation of the number of components with respect to *ϕ* under different follow probabilities *P*. Each curve corresponds to a fixed *P* ; points span all (*λ, r*) configurations mapping to a given *ϕ* = *λπr*^2^, and each value is the mean over 100 independent runs (error bars show the across-run standard deviation). As *ϕ* increases, all curves decrease and gradually stabilize. The curves corresponding to different values of *P* are clearly separated, and the theoretical predictions agree well with the simulation results across the entire range, confirming the model’s validity for the geometric density parameter *ϕ*.

Therefore, *η* jointly characterizes the effective interaction strength governing component formation.

Using *η* as the horizontal axis collapses the combined effects of spatial density and behavioral following tendency into a single control parameter. The theoretical prediction of the model and the simulation results are overall consistent, both showing a decreasing trend as *η* increases. Figure 5B shows a comparison between model and simulation results over *ϕ* = *λπr*^2^, the dimensionless geometric density parameter, for different follow probabilities *P* ∈ {0.5, 0.6, …, 0.9}. As *ϕ* increases, we see a decrease followed by saturation. This seems intuitive as higher agent densities result in fewer components. The model predictions agree well with the simulation results over the entire range, confirming the model’s predictive power.

The empirical term in eq. (6) is the geometric correction function *K*(*λ, r*). Its coefficients are calibrated by weighted least squares on the 20% of conditions covering extreme values of *λ* and *r*. The model achieved a mean absolute percentage error (MAPE) of 0.016 and a root-mean-square error (RMSE) of 0.84 on the validation set. Errors were computed separately for each parameter configuration, taking the mean over 100 repeated simulations, and then averaged across all validation conditions. The dataset was partitioned into 260 calibration conditions and 1040 validation conditions. To quantify uncertainty, we constructed approximate 95% prediction intervals assuming Gaussian fluctuations around the predicted mean. For each parameter configuration, the interval width combined the empirical standard error estimated from repeated simulations with an additional residual variance term estimated from the calibration set. The resulting empirical coverage rate on the validation set was 99.2%, indicating that the theoretical model accurately captures both the mean trend and the variability of the observed component statistics across a broad parameter range. The empirical coverage slightly exceeds the nominal 95% level, suggesting that the prediction intervals are mildly conservative due to the inclusion of both simulation uncertainty and residual model variance.

### 5.2 Simulation experiments

To showcase the applicability of our framework to engineering problems in distributed systems, we studied a simple aggregation scenario for a multi-robot system (see Appendix 7.3). We evaluate the controller on 42 simulated robots (Fig. 6), all with desired group size *G*^*^ = 14, over 1000 independent runs per signaling mode.

**Figure 6:**
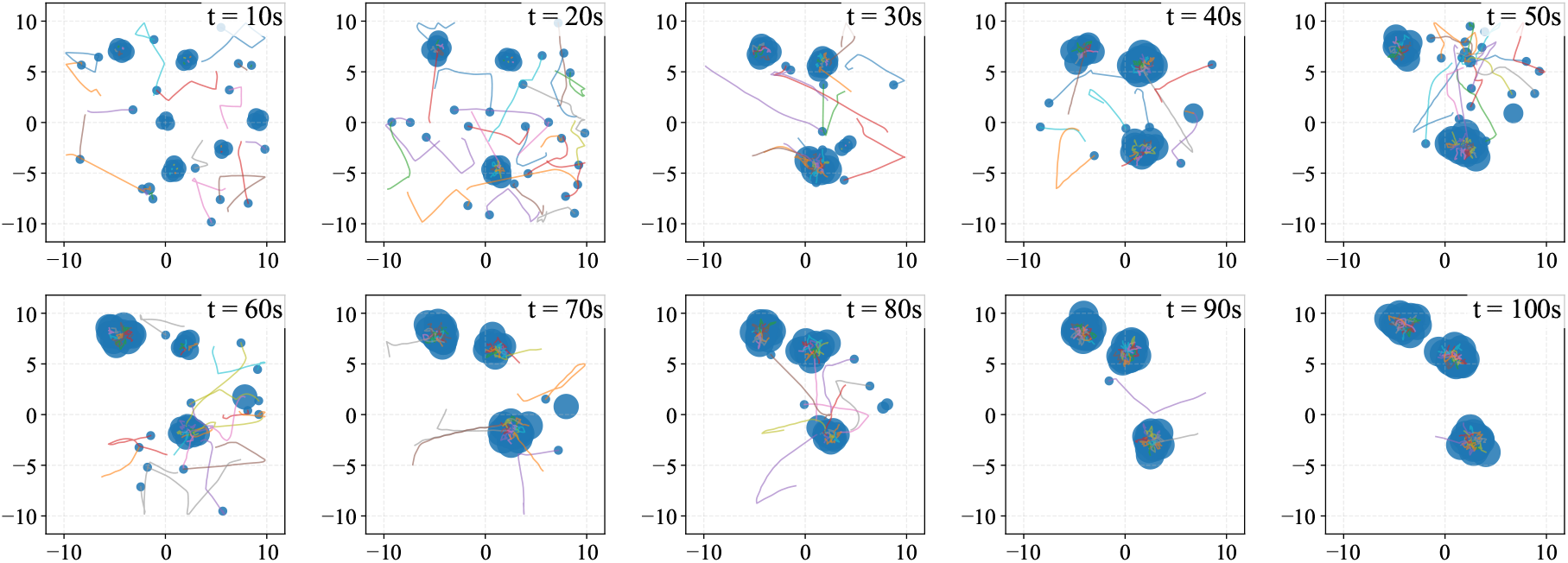
Snapshots of a representative simulation run with 42 robots. Robots determine their duration within a group based on the group’s current size (see Figs. 3A and 4 for the underlying decision mechanism). When the group is small and no new members join, robots leave and search for other groups; when a group becomes too large, robots probabilistically switch to fission behavior and leave the group. With the target group size of *G*^*^ = 14, the system continuously experienced the formation and dissolution of small groups, along with dynamic adjustments within larger groups, throughout the simulation. After 100 seconds, all robots have aggregated into one of three groups, each consisting of 14 individuals, thereby achieving the objective.

#### 5.2.1 Temporal availability of group-size information

In our framework, agents may obtain group-size information from neighboring groups before physically joining them. Group sizes are estimated in a decentralized manner using a round-based extension of extrema propagation Baquero et al. (2011) that runs continuously, allowing estimates to track membership changes under fission-fusion dynamics.^1^ Different signaling methods provide group-size information with different temporal availability during the joining process.

We compare three signaling and decision modes. In position-only signaling, agents select targets only based on positional information from nearby neighbors without explicit group-size information. In one-shot group-size signaling, agents use the communicated group size only once when selecting a follow target, and afterward continue following based only on positional information, without switching targets. In continuous group-size signaling, agents continuously receive updated group-size information and may revise their follow decisions over time. These signaling regimes allow us to study how the temporal availability of group-size information influences decentralized group regulation.

#### 5.2.2 Stability

We find that each of the three signaling and decision modes shows different group dynamics. In the position-only mode, groups tend to oscillate because, although oversized groups shed members through fission, larger groups still attract new members more frequently since random target selection is performed over individuals rather than groups, making members of larger groups more likely to be selected. The one-shot signaling mode reduces the amplitude of oscillations, but local instabilities still occur; that is, groups may have exceeded desired sizes by the time additional robots join. The continuous signaling and decision mode exhibits the best stability, eliminating oscillations through feedback and enabling the robot to accurately track changes in the target swarm size.

To quantify stability differences, we use *oversize time* (i.e., the time a robot spends in a group larger than the desired size) as the metric, defined as the cumulative duration during which a robot’s group size exceeds *G*^*^ = 14, averaged across all robots in a trial. This metric reflects the asymmetric oscillation mechanism described above: since larger groups contain more individuals, robots selecting a random target are more likely to join them, causing the system to dwell above *G*^*^ rather than below it, making oversize time a natural measure of instability in this context. The results (see Fig. 7A) show that: in the position-only mode, the system shows oscillatory characteristics, with an average oversize time of 47.38 s (± 14.71 s), which reflects the limitation of relying only on local sensing. The introduction of one-shot signaling significantly reduces the oversize time to 8.13 s (± 7.53 s), with a reduction of 82.8 %. This confirms that access to group-size information is the key driver of the improvement: even a single query at the moment of target selection is sufficient to substantially enhance system performance. The continuous signaling and decision mode further limits the oversize time to a narrow range of 2.18 s (± 2.49 s), a reduction of 95.4 % relative to the position-only baseline.

**Figure 7:**
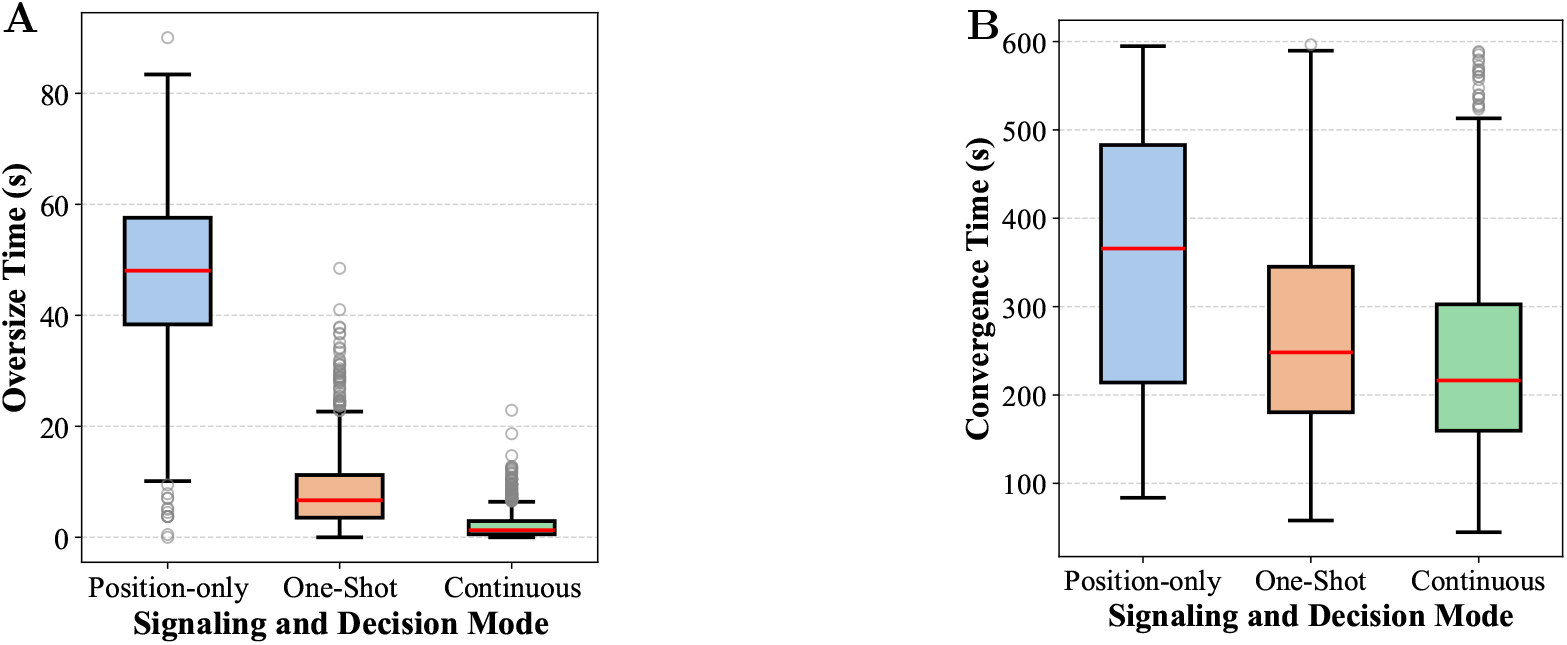
Performance comparison of different signaling and decision strategies. (A) Oversize time (period of time during which the size of the group exceeds the desired value) under varying signaling and decision levels. As the degree of communication increases, both median oversize time and oscillation decrease significantly. Continuous signaling and decision strategy nearly eliminates the oversize phenomenon, whereas position-only signaling and decision strategy leads to the most severe and unstable oversize. (B) Convergence time under the same settings. Communication markedly accelerates convergence compared with the position-only signaling and decision strategy, confirming its stabilizing effect on group coordination.

#### 5.2.3 Convergence time

We define convergence time as the duration from the start of the experiment until all robots have reached and stably maintained the desired group size *G*^*^ = 14. Trials in which any robot fails to reach *G*^*^ within the 1200s window are recorded as non-converged and excluded from the convergence time statistics but counted in the success rate. The results show that the three signaling and decision modes differ significantly in system convergence performance (see Fig. 7B). The continuous signaling and decision mode performs optimally with an average convergence time of 239.15 s (standard deviation 108.21 s) and a success rate of 91.9% within 1200 s. The one-shot signaling mode was the next best, with an extended average convergence time of 271.13 s (standard deviation 119.91 s) and a success rate of 84.4%. The position-only mode, on the other hand, was the least efficient, with an average convergence time of 357.49 s (standard deviation 140.66 s) and a success rate of only 10.9%.

These data show that making group-size information available to the agents improves system performance: continuous signaling and decision mode reduce the convergence time by 33.1% and increase the success rate by a factor of 8.4 compared to position-only signaling. Notably, even with a one-shot signaling mode, a success rate of more than 80% is achieved, providing an important reference for application scenarios that need to balance communication overhead and system performance.

### 5.3 Empirical case study using coati tracking data

To examine whether the proposed controller can generate aggregate patterns comparable to those observed in a natural fission–fusion system, we conducted an empirical case study using tracking data on the white-nosed coati (*Nasua narica*) from Grout et al. Grout et al. (2024).

As noted in Sec. 4, it is important to clarify the distinction between the calibration phase and the runtime behavior of the controller. The subjective group-size preferences 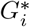 are inferred offline from empirical tracking data prior to any simulation. Each agent’s preference is set once, as a fixed scalar, before the simulation begins. The calibration step grounds the individual-level parameters in biological observations, but does not introduce a global feedback channel at runtime.

We calibrate group-size preferences from empirical subgroup-size summaries and then compare selected simulated and observed statistics. They conducted full-colony GPS tracking of white-nosed coati groups in the tropical forests of Panama, capturing individuals’ locations and tracking changes in group dynamics under natural conditions.

The resulting group-size and group-number distributions serve as consistency checks. The in-sample consistency check is whether the simulated transition dynamics between group-number states, a temporal pattern not encoded in 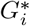, match the empirical Markov structure rather than a random baseline.

#### 5.3.1 The coati fission–fusion system

White-nosed coatis provide a particularly suitable biological system for evaluating the proposed framework because their fission–fusion dynamics are believed to be shaped by social organization rather than by external ecological constraints. Female adults and juveniles form relatively stable social bands, whereas adult males typically remain peripheral or solitary. As a result, subgroup formation in coatis appears to involve persistent social preferences and relatively stable tendencies in the size of subgroup associations.

#### 5.3.2 Biological rationale for the modeling assumptions

This structure aligns closely with the central assumption of our model, that individuals possess subjective group-size preferences and generate behavioral responses based on perceived local group size. Environmentally driven fission–fusion dynamics, for instance those shaped by resource distribution or predation pressure in shoaling fish or ungulate herds Kelley et al. (2011); Della Libera et al. (2023), can in principle be accommodated within the same framework by letting the subjective preferred group size *G*^*^ depend on these ecological variables. Agent-based models have demonstrated, for example, that simple foraging decisions in heterogeneous resource landscapes can generate complex fission–fusion organization Ramos-Fernández et al. (2006). Rather than requiring a distinct mechanism, such cases correspond to a preference that is itself modulated by external conditions, a natural extension we leave to future work.

#### 5.3.3 Data and calibration

Grout et al. Grout et al. (2024) tracked three groups, of which two exhibited fission–fusion dynamics (*Presidente* and *Galaxy*). The third group (*Trago*) remained cohesive throughout the observation period and was also excluded from the subgrouping analyses in that study. The raw data consists of position coordinates of all group members during daytime active periods, with a recording interval of 10 minutes. For the *Presidente* group, we use all *N* = 22 tracked individuals available in the dataset over the window in which they have simultaneous data. This exceeds the core group size of 16 reported in their main text, as we also retain peripheral individuals (e.g., largely solitary adult males) whom they excluded from group-size counts, since these, too, are agents with their own group-size preferences in our model. Subgroups were identified at each 10-minute sample by clustering GPS positions (DBSCAN,

50 m, following Grout et al. Grout et al. (2024)). For *Galaxy*, which has regular and symmetric splitting, we use a single common preference; for *Presidente*, each agent receives its own rounded mean experienced subgroup size 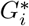 (Table 1). Simulations use the one-shot signaling mode; see Appendix 7.4.1 for details.

**Table 1:** Subjective group-size preferences for the two coati groups and their empirical basis. Values are grounded in the subgrouping structure reported by Grout et al. Grout et al. (2024). *Galaxy* splits consistently into two stable subgroups of comparable size, motivating a single common preference; *Presidente* splits unevenly and is far more heterogeneous, motivating per-individual preferences.

|  | *Galaxy* | *Presidente* |
| --- | --- | --- |
| Tracked individuals ($N$) | 11 | 22 |
| Observed subgrouping | two stable subgroups<br>(about 5 + about 5) | uneven split<br>(about 12 + about 4) |
| Experienced group size | concentrated (about 7) | heterogeneous (1 to 13) |
| Preference $G^*$ | common, $G^* = 6$ | per-individual rounded mean |
| Tolerance | 5 (range [1, 11]) | — |
| Basis | group structure | individual means (Fig. 8) |

#### 5.3.4 Reproducing empirical group-size and group-number patterns

We input the subjective group-size preferences 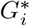 extracted from real coati data into each agent’s decentralized fission–fusion controller. After calibration, the simulations produced group-size and subgroup-count summaries that were similar to several features of the empirical data, while retaining quantitative differences (see Fig. 8, middle and bottom rows). We compare the central tendencies and frequencies descriptively rather than treating their agreement as an independent model prediction. In both *Galaxy* and *Presidente* groups, the simulated distributions closely matched the real data, capturing not only the central tendency of group sizes but also the variability across individuals and the frequency of group numbers. For the *Galaxy* group, the mean group size for all individuals in the real data was approximately 7.2 (±0.5), whereas the simulation yielded a comparable mean of 8.0 (±0.5). The distribution of group numbers showed that both real and simulated data were dominated by group numbers one, two, and three subgroups, with state two being the most frequent (42.1% in real data vs. 47.9% in simulation), followed by state three (28.1% vs. 25.0%) and state one (21.7% vs. 25.0%). Higher-order states (four and five) were rare in both real and simulated systems, indicating similarity in these selected marginal subgroup-count frequencies in the subgroup structure of *Galaxy* after calibration. In the *Presidente* group, real data indicated a much larger mean subgroup size of 9.7 (± 4.4), whereas the simulation generated a slightly smaller but comparable mean of 8.5 (± 2.8). The subgroup number distribution was broader than in *Galaxy*, with states four, five, and six dominating the dynamics. In particular, states five and six accounted for more than half of the time in the real data (28.1% and 28.1%), closely matching the simulation (26.1% and 21.9%). State four was also frequent (17.3% vs. 17.9%), and higher subgroup numbers (between seven and nine) appeared occasionally in both real and simulated data, though at low frequency. These results showcase that our controller approach can reproduce the contrasting patterns of fission–fusion dynamics across groups: frequent binary fissions in *Galaxy* versus larger, more stable multi-subgroup structures in *Presidente*.

**Figure 8:**
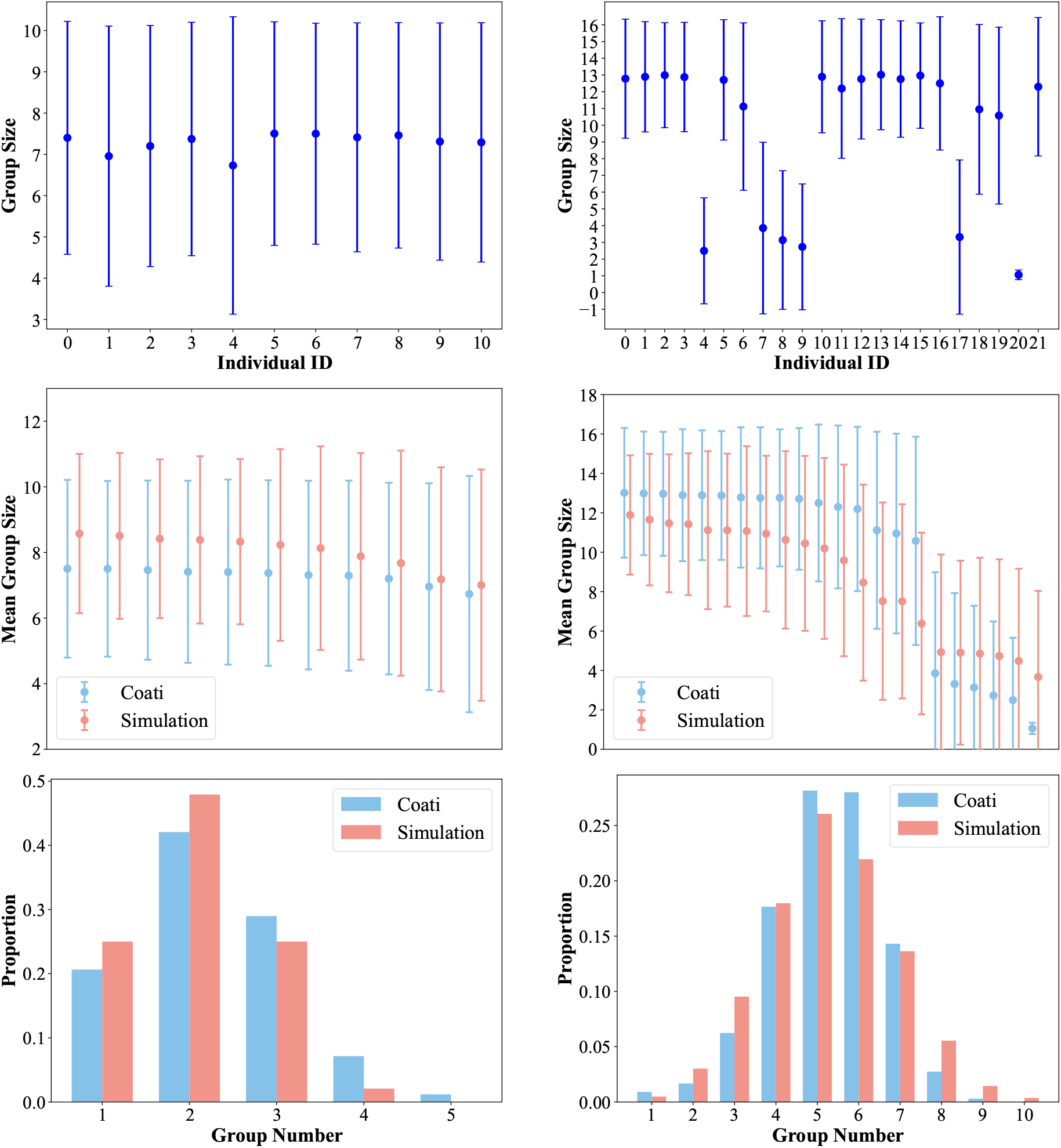
Comparative analysis of group size characteristics in *Galaxy* (left column) and *Presidente* (right column). Top row: Mean group size estimated from GPS tracking data of coatis, obtained by clustering individuals at each time step and calculating the mean and standard deviation of their group sizes. Middle row: Comparison between controller-based simulation results and empirical coati data with respect to mean group size, ranked in descending order. Bottom row: Comparison of the distributions of group numbers per time step between simulated data and empirical observations.

#### 5.3.5 Reproducing empirical fission–fusion transition dynamics

We summarize the dynamics by the subgroup count *G*_*t*_ obtained from the same clustering. We remove consecutive repeated observations before counting transitions, so the estimated matrices are embedded (jump) chains with zero diagonal. We use a Gaussian-weighted matrix as a data-independent reference, in which the probability of moving from *i* to *j* decays with (*i*− *j*)^2^ (*σ* = 1). Matrices are compared by the row-averaged Jensen–Shannon distance Lin (1991) (Appendix 7.4.2).

As shown in Fig. 9 top and middle rows, both empirical and simulated jump-chain matrices exhibited predominantly local changes in subgroup count, with transitions concentrated between nearby states. These transitions concentrated between neighboring subgroup counts (e.g., 2 ↔ 3, 3 ↔ 4). This indicates that changes in the number of subgroups were typically gradual rather than involving large jumps. By contrast, the Gaussian-weighted baseline (Fig. 9, bottom row) captured only locality in the state space and lacked the characteristic transition structure observed in the empirical and simulated data. Consistent with this qualitative agreement, the Jensen–Shannon distance between the empirical and simulated matrices was low for both groups (*Galaxy*: 0.030; *Presidente*: 0.148), whereas the Gaussian baseline was substantially more distant (*Galaxy*: 0.240; *Presidente*: 0.309). These results confirm that the proposed controller reproduces key aspects of the fission-fusion transition dynamics observed in wild coatis, beyond what a simple, data-independent baseline captures.

**Figure 9:**
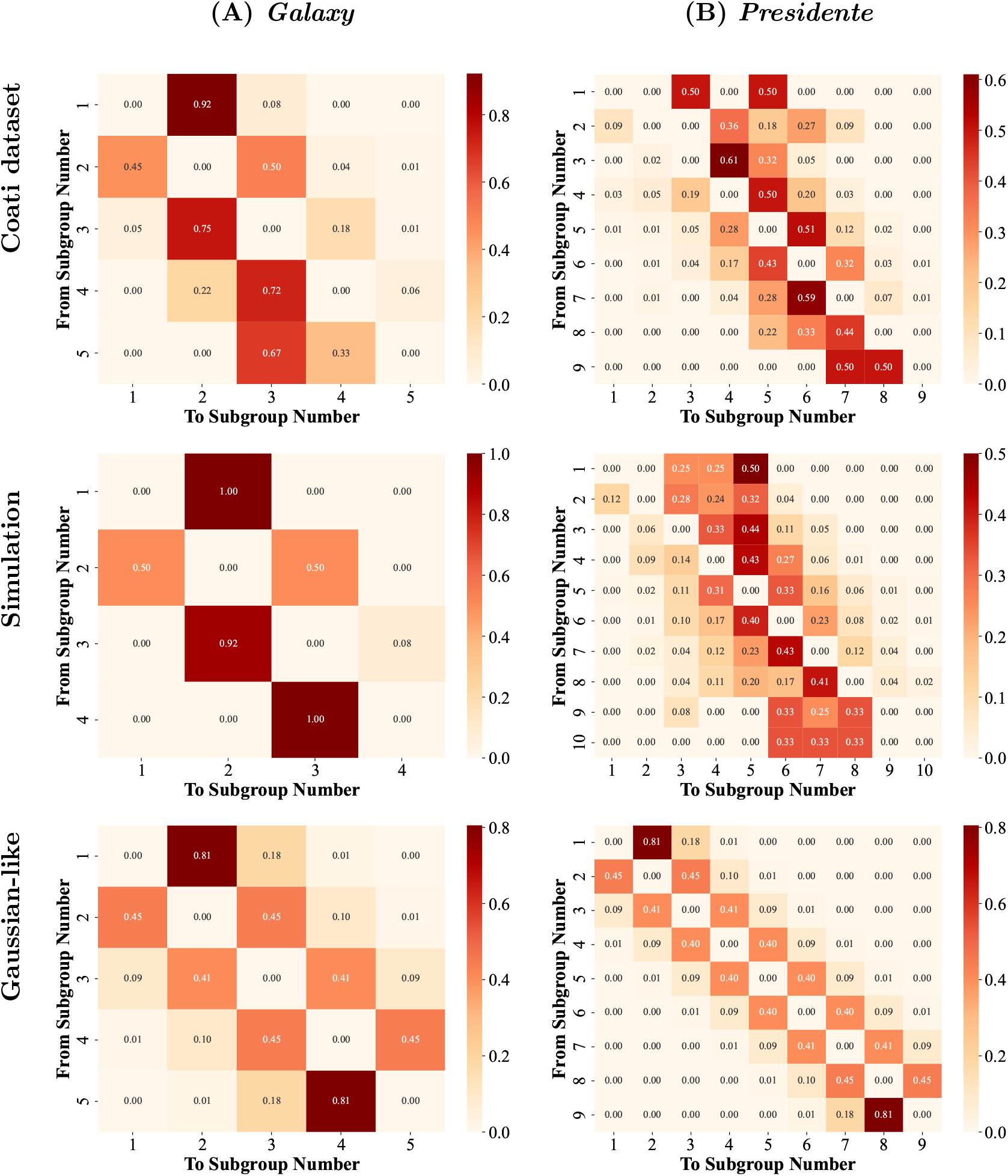
Markov transition analysis of subgroup count dynamics for *Galaxy* (left column) and *Presidente* (right column). Top row: Transition matrices derived from GPS tracking data of coatis, representing the empirical dynamics of subgroup number changes. Middle row: Transition matrices generated by the controller, in which real group size data were used as input states. Bottom row: Baseline transition matrices randomly generated from a Gaussian distribution with the same dimensionality as the empirical data, serving as a data-independent locality baseline. The comparison of calibrated simulations with selected empirical statistics, relative to the random baseline.

## 6 Conclusion

We introduced the Group Size and Number Regulation Problem (GSNRP) to ask whether locally interacting individuals can jointly regulate the sizes and number of groups without explicit coordination. The framework builds on the established size-biased sampling relationship between group-level and individual-level descriptions of group size Jarman (1974); Reiczigel et al. (2008); Jovani and Mavor (2011). For a population partitioned into groups,

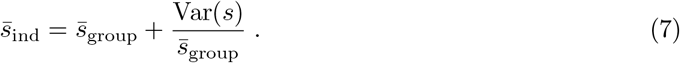

Thus, for a fixed population size, group number or mean group size alone is insufficient to characterize the social conditions experienced by individuals whenever group sizes are heterogeneous. A mechanistic account of social organization must therefore consider the group-size distribution and group number as coupled outcomes.

Prior work has provided complementary accounts of group-size optimality, split–merge dynamics, and emergent group-size distributions. Fitness-based models search for an optimal group size Sibly (1983); Brown (1982); Markham et al. (2015). In contrast, population-level models characterize group-size distributions Bonabeau et al. (1999); Niwa (2003). The connection between these perspectives is less often formulated explicitly as a decentralized feedback-regulation problem. The GSNRP builds on the size-biased sampling relationship to distinguish group-level and individual-level descriptions (Definition 1): The individual-level mean 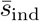 systematically exceeds the group-level mean 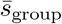 whenever the size distribution has nonzero variance, because individuals experience groups in proportion to group size. Consequently, any mechanism that regulates social organization must jointly account for both quantities rather than treating them independently.

The minimal follow model (Sec. 3) demonstrates that local interaction rules alone are sufficient to produce group structures. In our framework, group-size distributions are not prescribed but emerge from repeated individual decisions to split or merge in pursuit of a preferred group size.

The minimal following model showed that simple local interactions generate predictable group structures. The resulting graph-theoretical approximation accurately predicted the number of components across held-out parameter conditions. However, because the resulting structure is determined by density, interaction range, and the following probability, the model provides no feedback to maintain groups near a preferred size. This limitation motivated the introduction of subjective group-size preferences and decentralized fission–fusion responses.

Our proposed fission-fusion mechanism goes beyond reproducing group-size distributions: it provides a decentralized mechanism by which these distributions Bonabeau et al. (1999); Niwa (2003) emerge as a consequence of individuals pursuing their preferred group size, where this preference may reflect the fitness consequences of group living Brown (1982); Markham et al. (2015). The observed population-level distribution is the aggregate outcome of repeated individual decisions to split or merge.

The robot simulations showed that this feedback mechanism can regulate group structure and that its performance depends strongly on the availability of group-size information. The three signaling modes reveal that the availability of group size information is the critical factor for stable regulation: even one-shot signaling reduces oversize time from 47.38 ± 14.71 s to 8.13 ± 7.53 s. The biological validation with coati data was also conducted under one-shot signaling, and the controller nevertheless reproduced the empirical fission–fusion dynamics of both groups, supporting the biological plausibility of the proposed mechanism.

The simulations show that decentralized size-dependent decisions are sufficient to generate similar patterns within the model. While they do not demonstrate that real coatis base their fission–fusion decisions on group size, the results are consistent with the hypothesis that animals continuously regulate their social environment through repeated decisions to split or merge in response to local conditions.

Our framework assumes that individuals can assess their current group size, at least approximately. Direct evidence for such an ability is limited. Although an exact count may not be necessary in principle, the mechanism could instead operate on a signal that varies monotonically with local membership and can be compared with a reference level. Whether such indirect cues are sufficient was not tested here. Identifying plausible proxies for this signal is a natural next step.

Future work should test the mechanism on additional populations and time periods, compare it with alternative behavioral models, examine robustness to noisy or indirect group-size cues, and allow preferred group size to vary with ecological conditions or task demands.

In summary, a simple decentralized feedback mechanism based on local group-size estimates can, without global coordination, generate selected group-size and group-number distributions resembling those observed in wild coatis after calibration. The controller provides a compact phenomenological model and a testable hypothesis for how individual preferences might shape collective social organization.

## Data availability

The coati tracking data are openly available at https://doi.org/10.5281/zenodo.11204635 Grout et al. (2024).

The controller source code and ARGoS configuration files are available at https://github.com/Tianfu-swarm/fission_fusion_controller. The generated simulation data are archived at Zenodo: https://doi.org/10.5281/zenodo.22070402 Zhang et al. (2026).

## Author contributions

T.Z.: conceptualization, methodology, software, formal analysis, investigation, validation, visualization, writing—original draft; S.L.: conceptualization, visualization, writing—review and editing; H.H.: conceptualization, methodology, supervision, project administration, funding acquisition, writing—review and editing. All authors gave final approval for publication and agreed to be held accountable for the work performed therein.

## 7 Appendix

### 7.1 Graph-theoretic model of group formation: derivation and calibration setup

We analyze the group structures generated by the minimal interaction model (see Sec. 3) from a graph-theoretic perspective. The goal is to characterize how group size and group number emerge from simple local interactions and how they depend on system parameters such as the interacting range within which an individual can select another individual as an interaction partner and the probability of choosing to follow.

#### 7.1.1 Graph-theoretic abstraction of the model

In terms of graph theory, the following relationships can be represented as a directed graph Γ = (*V, E*) with individuals as nodes *V* and the following behavior represented as directed edges *E* = {(*i, f* (*i*)) | *i* ∈ *V, f* (*i*) ≠ ⊥}. This type of graph is called a directed pseudoforest (outdegree at most one: ∀*i V* : deg^+^(*i*) ≤ 1) with each connected component being a directed pseudotree containing at most one cycle. Each pseudotree represents a group of individuals. There are two types of groups: those with a cycle and those without. Without a cycle (Fig. 3C), we get a directed tree that has a root *r* with *f* (*r*) = ⊥. With a cycle (Fig. 3D), a subset of nodes forms a directed cycle, and every remaining node has a directed path, following the map *f*, that eventually reaches this cycle.

#### 7.1.3 From local interaction range to expected rooted groups

We now embed the interaction model in physical space to relate local following interactions to global group structure. Consider *N* individuals distributed over a bounded two-dimensional area *A*. Placing individuals in a bounded area captures a basic ecological constraint built into the model: individuals can only follow nearby conspecifics. We define population density as

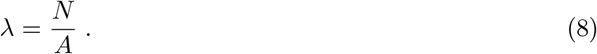

We further define the dimensionless geometric density parameter

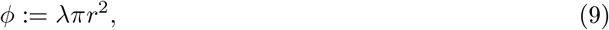

which corresponds to the expected number of neighbors within the interaction range *r* under the Poisson approximation. Intuitively, *ϕ* characterizes the system’s effective local connectivity and jointly captures the influence of population density and interaction range on group-formation dynamics.

We assume a random geometric graph. Let nodes *x*_1_, …, *x*_*N*_ ∈ *V* be sampled independently and uniformly from a bounded region *A* ⊂ ℝ^2^, that is, according to a binomial point process.

Edges are defined by range as defined above: (*i, j*) ∈ *E* ⇐⇒ ∥*x*_*i*_ − *x*_*j*_∥ ≤ *r* Penrose (2003). The probability that a node has at least one neighbor under a Poisson approximation is

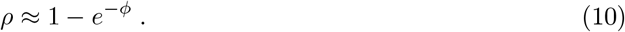

The probability that a node is the root is

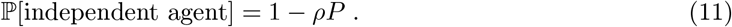

The expected number of directed pseudotrees in the pseudoforest is

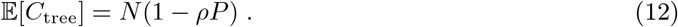

#### 7.1.3 Expected number of cyclic components

Cycle formation requires a closed chain of following decisions and is therefore more sensitive to spatial and behavioral constraints than the formation of rooted, cycle-free groups. Unlike cycle-free components, cyclic components can arise only when a sequence of local following decisions closes back on itself, making their expected number strongly dependent on both the spatial neighborhood structure and the probability of following.

In classical random mapping theory, the number of cyclic components can be obtained by normalizing the number of cyclic nodes by the average cycle length Flajolet and Odlyzko (1989). However, these results typically rely on the assumptions of a uniform node distribution and unbiased target selection. In our system, due to probabilistic following behavior, some nodes exhibit an out-degree of 0, thereby violating the original uniformity assumption by Flajolet and Odlyzko Flajolet and Odlyzko (1989). These constrained nodes no longer participate in the free connection process, thereby reducing the number of nodes that can form cycles. In addition, due to the limited follow range, connections are restricted to spatial neighbors, which reduces the effective connectivity among unconstrained nodes and decreases the number of nodes that participate in cycle formation.

To address these two constraints, we provide an approximate characterization of the number of cyclic components in a system subject to neighborhood and follower structure constraints. Our basic approach is to model and correct for these two types of influences while retaining the structural framework of classical random mapping results. First, by excluding nodes with zero out-degree, we derive the effective number of nodes participating in the mapping process, yielding *m*_eff_, and substitute this into the classical result to estimate the expected number of cyclic nodes. Second, we derive an effective average cycle length and incorporate correction factors to account for spatial non-uniformity. Finally, we convert the estimated number of cyclic nodes into the number of cyclic components by normalizing with the effective average cycle length, where the correction function *K*(*λ, r*) is calibrated experimentally in subsequent steps. A node with out-degree zero cannot contribute to cyclic structures. We define the effective follower probability as

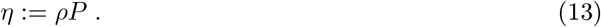

Accordingly, the number of nodes with positive out-degree is *N*_eff_ := *Nη*. A node can participate in a cycle formation only if it successfully follows (one factor *η*) and the node it follows also has positive out-degree, so that the following chain can continue (a second factor *η*). Hence the effective number of nodes entering the random mapping process is

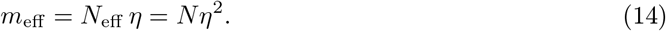

Substituting *n* → *m*_eff_ = *N*_eff_ *η* into the asymptotic result Flajolet and Odlyzko (1989), and introducing a geometric correction factor *ψ*(*λ, r*) ∈ (0, 1] to account for spatial non-uniformity arising from the finite follow range, the expected number of cyclic nodes is

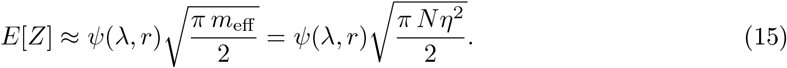

In the classical random mapping model with *n* nodes, the expected number of cyclic nodes scales as 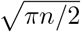 while the expected number of components scales as 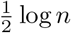 Flajolet and Odlyzko (1989).

Since each component contains exactly one cycle, the average cycle length is the ratio of these two quantities, 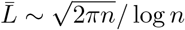 Flajolet and Odlyzko (1989). Substituting *n* → *m*_eff_ = *Nη*^2^ we get

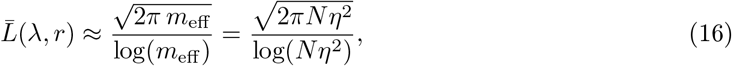

By incorporating the geometric correction factor, we get

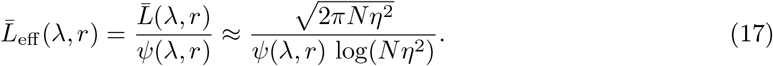

Dividing eq. (15) by eq. (17), the 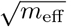 terms cancel, leaving a constant factor *ψ*^2^*/*2. We absorb this into a single correction function

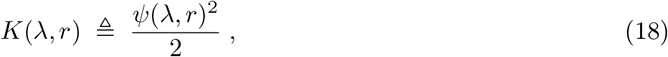

yielding the expected number of cyclic components:

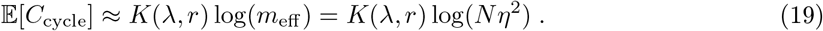

The parameters of *K*(*λ, r*) will be calibrated experimentally in subsequent steps, which will also give us an empirical validation.

#### 7.1.4 Expected total number of components

The total number of components is obtained by adding the tree-like and cyclic contributions. Combining eq. (12) and eq. (19), the expected total number of components is

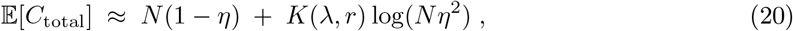

which is eq. (6) of the main text, where *η* = *ρP* and 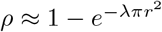.

#### 7.15 Simulation setup and calibration of K(λ, r)

To evaluate the predictive capability of the theoretical model for component structures derived above, this study conducted simulation experiments in a circular area with radius *R* = 10 m. The following parameters were set: follow probability *P* = {0.5, 0.6, …, 0.9}, follow range *r* = {1.0, 2.0, …, 10.0}, and the number of individuals *N* = {50, 60, …, 300}, to obtain the agent density *λ* = *N/A*. Each parameter combination was independently repeated 100 times, with the initial position and random seed generated at each repetition.

For each run, we generate a directed follow graph in which each node has at most one outgoing edge. Whether a node has an outgoing edge depends on two aspects: (1) the follow probability *P* and (2) the availability of neighbors within the perception radius *r*. If there is no neighbor, or if a probabilistic no-follow decision (with probability 1 −*P*) indicates not to follow, the node has zero out-degree. We use depth-first search (*DFS*) to identify components and record the relevant node and component information, calculating the total number of components *C*_total_, the number of tree components *C*_tree_, and the number of components containing cycles *C*_cycle_.

The theoretical model uses (*P, r, λ*) as independent variables and provides analytical expressions for the expected number of cyclic components, expected cycle length, and expected number of tree components as derived above. The only empirical term in the model that lacks a closed-form derivation is the correction function *K*(*λ, r*) = *ψ*(*λ, r*)^2^*/*2, where the geometric correction factor *ψ*(*λ, r*) ∈ (0, 1] accounts for the reduced cycle-formation probability under spatially constrained following. To calibrate *K*(*λ, r*), we select the 20% subset of the experimental data covering extreme parameters (maximum/minimum *λ* and maximum/minimum *r*) as the calibration set. Since the geometric correction factor *ψ*(*λ, r*) lacks a closed form, we estimate *K* empirically: for each condition we recover the empirical *K* from the observed cyclic-component count via eq. (19), and fit its coefficients by weighted least squares over the features [1, log(1 + *λπr*^2^), log(1 + *λπr*^2^)^2^, (*r/R*)^2^], with weights equal to the inverse across-repetition sample variance so that more consistent conditions contribute more to the fit. The estimated coefficients of *K*(*λ, r*) form a single shared set (not one factor per condition). These are then held fixed and applied to the remaining 80% of the conditions (the validation set), which therefore introduce no free parameters. We compared the deviation between the mean as predicted by theory eq. (20) and the observed mean, using the mean absolute percentage error (MAPE), the root mean square error (RMSE), and the empirical coverage of the prediction interval.

### 7.2 State definitions and transition logic of behavioral state-machine

#### 7.2.1 State Transition Conditions

The following three core elements determine state transitions in the behavioral state-machine model (see Sec. 4.1):

1. Subjective Group Size Preference: Each agent *i* possesses an individual desired group size 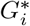 and estimates the size 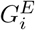 of the group it currently stays in. This desired size serves as an internal reference: when 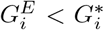 the agent perceives the group as too small and waits for new members. If the group remains undersized after a waiting period, the agent leaves via fission, resumes random exploration, and seeks to fuse with a new group. When 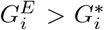, the agent perceives the group as too large and initiates fission with a certain probability, otherwise staying in the group. In both cases, the normalized deviation between 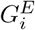 and 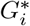 governs the strength of the response, as detailed in the following conditions.
2. Waiting Time: If the agent estimates that the current group size is smaller than its preferred size (*G*_*E*_ *< G*^*^), we call this a perceived group-size shortfall, and the agent activates a waiting behavior. We define the normalized perceived quantity below the optimal size as

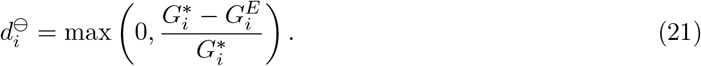 The waiting time decreases linearly with the perceived undershoot and is given by

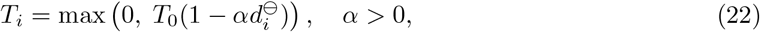

where *T*_0_ is a baseline waiting time corresponding to a near-optimal group size, and *α* (Fig 3E) controls the sensitivity (slope) of the waiting response. If the desired size *G*^*^ is not reached within time *T*, the agent enters the `Fission` state.
3. Follow Decision: The agent determines whether to follow a neighboring target based on the following range *r* and follow probability 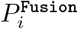. During `Random Walk`, agents explore the environment; upon detecting a neighbor within range *r*_random_, they transition to the Fusion state with a fixed probability *P*_random_. Otherwise, the agent transitions to the `Stay` state, forming a singleton group and waiting for others to join.

During `Stay`, the probability is adjusted dynamically based on the deviation from the optimal group size. We define the normalized perceived quantity above the optimal size as

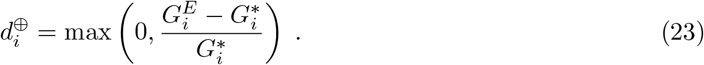

The follow probability is then given by

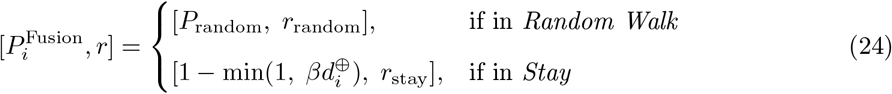

where *β >* 0 (Fig. 3E) is the slope parameter controlling the sensitivity of the response to oversized groups.

#### 7.2.2 State Functions and Transition Mechanism

As described above, we define the agent behavior microscopically as a state machine of four states: `Random_Walk`, `Fusion`, `Fission`, and `Stay` (Fig. 4).

1. `Random_Walk`: Agents perform a random walk by sampling linear and angular velocities independently from truncated Gaussian distributions. When a neighboring agent is detected within its follow range *r*_random_, the agent transitions to the `Fusion` state with probability *P*_random_. Otherwise, it transitions to the `Stay` state, forming a singleton group and waiting for others to join.
2. `Fusion`: After transitioning to `Fusion`, the agent moves toward the target (a random neighbor). When the agent gets close to the target (*< d*_fusion_), it transitions to `Stay`. If during movement the distance to the target exceeds the follow range, or the target location is lost, the individual reverts to `Random_Walk`.
3. `Stay`: In `Stay`, agents maintain distance from group members, via Lennard-Jones-style attraction-repulsion forces Gazi and Passino (2003), in order to preserve stable local structure. The attraction-repulsion forces allow each agent to maintain a balanced distance from its neighbors, preserving the coherence of the local group. Each agent continuously monitors the current size of its own group 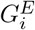 and calculates its own waiting time. When new agents join, its waiting time updates. If the agent’s desired group size is not reached by the end of the waiting time, the agent transitions to `Fission`.
4. `Fission`: When the current size 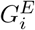 of the group to which an agent belongs is greater than its desired size 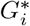, individuals enter `Fission` from `Stay` with probability 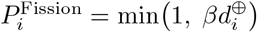. In the Fission state, individuals decide to separate and move away from their nearest neighbor in this group. Once the distance to their neighbors from the previous group exceeds *d*_fission_, they switch to *random walk*.

### 7.3 Simulation experiments setup and data

#### 7.3.1 Experimental platform and controller

To demonstrate the applicability of our framework to engineering problems in distributed systems, we examined a simple aggregation scenario in a multi-robot system. For the simulation study, we used the ARGoS simulation platform Pinciroli et al. (2012) to evaluate the effectiveness of swarm control strategies. We constructed a 20 *m* 20 *m* × arena with 42 simulated robots based on the FootBot Dorigo et al. (2013) platform, which served here as a convenient generic mobile robot platform rather than as a focus of the study. The communication range was set to 10 meters. The simulated robot platform implements proximity sensing, and the simulator runs a physics engine for collision detection.

#### 7.3.2 Experimental data

All robots were set to a desired group size of *G*^*^ = 14 members. First, we allow the robots to move in an initial five-second random-wandering phase to eliminate positional correlations. During the subsequent 1200 seconds of experimentation, we record: timestamp, internal ID of each robot, global position, its current group size, and communication events for the three signaling and decision modes. See snapshots of a representative simulation run in Fig. 6. To ensure statistical significance, 1000 independent repetitions of the experiment were performed for each signaling mode, resulting in a total of 3000 time-series datasets.

### 7.4 Coati data processing and analysis

#### 7.4.1 Calibration of Subjective Group-Size Preferences

For the empirical case study (see Sec. 55.3), we calibrate and validate the coati group-size dynamics. For each time interval, we identified the sub-group to which each individual belonged using a spatial clustering method (DBSCAN, with a radius threshold of 50 m, following Grout et al. Grout et al. (2024)). This clustering allowed us to estimate each individual’s experienced mean group size and associated variability (Fig. 8, top row), which we interpreted, for the sake of simplicity in this study, as the individual’s subjective group size preference. While this may be an ad hoc modeling assumption, it allows us to calibrate the model using parameters inferred from biological observations.

The subjective group-size preferences for the two groups are derived directly from the empirical subgrouping structure reported by Grout et al. Grout et al. (2024), and are summarized in Table 1. The two groups exhibit different subgrouping patterns, which we specify through preferences at different levels. The *Galaxy* group splits consistently into two stable subgroups of comparable size (about five individuals each), alternating between this even split and the intact group of roughly eleven Grout et al. (2024). This regular, symmetric structure is well captured by a single common preference *G*^*^ = 6, which is about half the full group, with a tolerance of five so that the admissible range [1, 11] covers both the split subgroups and the intact group. The *Presidente* group, by contrast, splits unevenly into a large (≈12) and a small (≈4) subgroup Grout et al. (2024), and its individuals differ widely in their typical subgroup sizes (per-individual means ranging from roughly one to thirteen; Fig. 8). Because these preferences are heterogeneous across individuals, we assign each agent its own empirical mean subgroup size 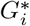, which is rounded to the nearest integer.

#### 7.4.2 Markov modeling of group-number dynamics

Let *G*_*t*_ ∈ {1, …, *G*_max_} be the number of subgroups at time *t*, where *G*_max_ is the largest subgroup count observed. The chain is characterized by a transition matrix **P** = [*p*_*ij*_] with *p*_*ij*_ = P[*G*_*t*+1_ = *j* | *G*_*t*_ = *i*] and Σ _*j*_ *p*_*ij*_ = 1. For the coati data, we cluster the GPS positions at each time step (DBSCAN, as above) to obtain a state sequence. Since the duration of a given subgroup count is not of interest here, we remove consecutive repeated observations, yielding the embedded (jump) chain. We count all state-to-state transitions *c*_*ij*_, and row normalize to obtain

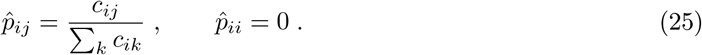

The transition matrix for the simulation experiments is estimated identically by clustering the simulated agents’ positions at each time step and applying the same state-sequence construction, transition counting, and row normalization. As a data-independent reference over the same state space, we construct a Gaussian-weighted baseline transition matrix

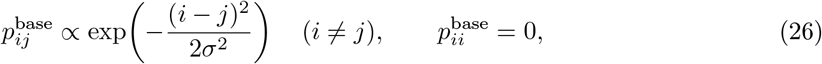

with *σ* = 1, followed by row-normalization. This concentrates weight on neighboring states but carries no data-specific information, yielding a locally structured reference that is independent of the observed data and does not encode any biological dynamics. We quantify the discrepancy between two transition matrices using the Jensen-Shannon distance Lin (1991). The distance is computed independently for each pair of corresponding rows using only entries that have positive probability in at least one of the two rows, and the resulting row-wise distances are averaged. The Jensen–Shannon distance is symmetric, bounded in [0, 1], and well-defined for matrices containing zero-probability entries.

## Footnotes

1 Implementation: https://github.com/Tianfu-swarm/fission_fusion_controller

